# CCR2-dependent monocyte-derived cells restrict SARS-CoV-2 infection

**DOI:** 10.1101/2021.05.03.442538

**Authors:** Abigail Vanderheiden, Jeronay Thomas, Allison L. Soung, Meredith E. Davis-Gardner, Katharine Floyd, Fengzhi Jin, David A. Cowan, Kathryn Pellegrini, Adrian Creanga, Amarendra Pegu, Alexandrine Derrien-Colemyn, Pei-Yong Shi, Arash Grakoui, Robyn S. Klein, Steven E. Bosinger, Jacob E. Kohlmeier, Vineet D. Menachery, Mehul S. Suthar

## Abstract

SARS-CoV-2 has caused a historic pandemic of respiratory disease (COVID-19) and current evidence suggests severe disease is associated with dysregulated immunity within the respiratory tract. However, the innate immune mechanisms that mediate protection during COVID-19 are not well defined. Here we characterize a mouse model of SARS-CoV-2 infection and find that early CCR2-dependent infiltration of monocytes restricts viral burden in the lung. We find that a recently developed mouse-adapted MA-SARS-CoV-2 strain, as well as the emerging B. 1.351 variant, trigger an inflammatory response in the lung characterized by expression of pro-inflammatory cytokines and interferon-stimulated genes. scRNA-seq analysis of lung homogenates identified a hyper-inflammatory monocyte profile. Using intravital antibody labeling, we demonstrate that MA-SARS-CoV-2 infection leads to increases in circulating monocytes and an influx of CD45+ cells into the lung parenchyma that is dominated by monocyte-derived cells. We utilize this model to demonstrate that mechanistically, CCR2 signaling promotes infiltration of classical monocytes into the lung and expansion of monocyte-derived cells. Parenchymal monocyte-derived cells appear to play a protective role against MA-SARS-CoV-2, as mice lacking CCR2 showed higher viral loads in the lungs, increased lung viral dissemination, and elevated inflammatory cytokine responses. These studies have identified a CCR2-monocyte axis that is critical for promoting viral control and restricting inflammation within the respiratory tract during SARS-CoV-2 infection.

## Main

Severe acute respiratory syndrome coronavirus 2 (SARS-CoV-2) is a novel β-coronavirus which emerged in Wuhan, China in December 2019 and is the causative agent of coronavirus disease 2019 (COVID-19)^1,2^. Innate immunity to SARS-CoV-2 begins with a limited interferon (IFN) response and production of inflammatory cytokines (IL-6, Il-1β, TNFα, IL-8) by respiratory epithelial cells or alveolar macrophages^3–6,7^. As shown in bronchoalveolar lavages (BAL) of COVID-19 patients, this innate immune response coincides with robust infiltration of neutrophils, monocytes, and dendritic cells into the lung airways^6,8^. Monocytes in the lung parenchyma can be divided into subpopulations characterized by their expression of Ly6C; Ly6C high classical monocytes are pro-inflammatory, whereas Ly6C low non-classical monocytes promote wound healing^9,10^. Ly6C low monocytes are prevalent during homeostatic conditions, however after a viral infection Ly6C high monocytes will infiltrate the lung in a CCR2 dependent manner^10–12^. Classical Ly6C high monocytes can differentiate into monocyte-derived dendritic cells (moDCs), which increase in number in response to viral respiratory infection, produce Type I IFN, and excel at antigen presentation^13^. The contribution of monocytes to promote protective immunity to SARS-CoV-2 infection is not known. In this study, we utilize a mouse-adapted (MA) SARS-CoV-2 strain and the human variant B.1.351 to evaluate the contribution of monocytes to protective immunity against SARS-CoV-2 and identify a CCR2-monocyte axis that is critical for promoting viral control and restricting inflammation within the respiratory tract during SARS-CoV-2 infection.

### Variant B.1.351 and MA-SARS-CoV-2 replicate in the lungs of C57Bl/6 mice

To investigate the immunological response to SARS-CoV-2 in the lung, we generated a MA-SARS-CoV-2 strain (Supplemental Fig. 1A). We engineered mutations into the icSARS-CoV-2 backbone^14^ that have been shown to increase SARS-CoV-2 virulence in mice^15^. Next, this virus was serially passaged 20 times in the lungs of Balb/c mice. Deep sequencing of plaque isolated virus revealed three additional acquired mutations which include two within the Spike (K417N and H655Y) and one within the Envelope (E8V) gene. Next, we confirmed the utility of MA-SARS-CoV-2 as a model to study SARS-CoV-2 pathogenesis in C57Bl/6 mice. Intranasally infected mice survived infection with MA-SARS-CoV-2, but had 10% body weight loss at day 2-3 post infection (p.i.) (Fig. 1A). Lung tissue was harvested at 0, 2, and 4 days p.i., and infectious MA-SARS-CoV-2 was measured via plaque assay. MA-SARS-CoV-2 titers peaked at day 2 p.i, with 10^9^ plaque forming units (PFU) per gram of lung tissue. Viral RNA peaked in the lung at day 2 p.i., dropping 100-fold by day 4 p.i (Fig. 1B). To determine localization of MA-SARS-CoV-2 in the lung, we performed *in situ* hybridization using probes that target the Spike gene of SARS-CoV-2. Viral RNA was restricted to cells lining the airways of the lung, in accordance with observations in humans (Fig. 1C)^16^. Examination of the antiviral response to MA-SARS-CoV-2 in the lung found expression of *Ifnl2* and ISGs peaked at day 2 p.i. Chemokines (*Cxcl10, Ccl2, Ccl5*) and endogenous pyrogens (*Il6, Tnf, Il1b*) were upregulated in response to MA-SARS-CoV-2 infection in the lung (Supplemental Fig. 1B). We next plotted gene expression of representative transcripts against viral RNA and found that *Ifih1, Ifnl2*, and *Il6* levels positively correlated with MA-SARS-CoV-2 viral burden (Fig. 1D). Thus, MA-SARS-CoV-2 infects the respiratory tract and induces a viral load dependent inflammatory response in C57Bl/6 mice.

**Figure 1.**
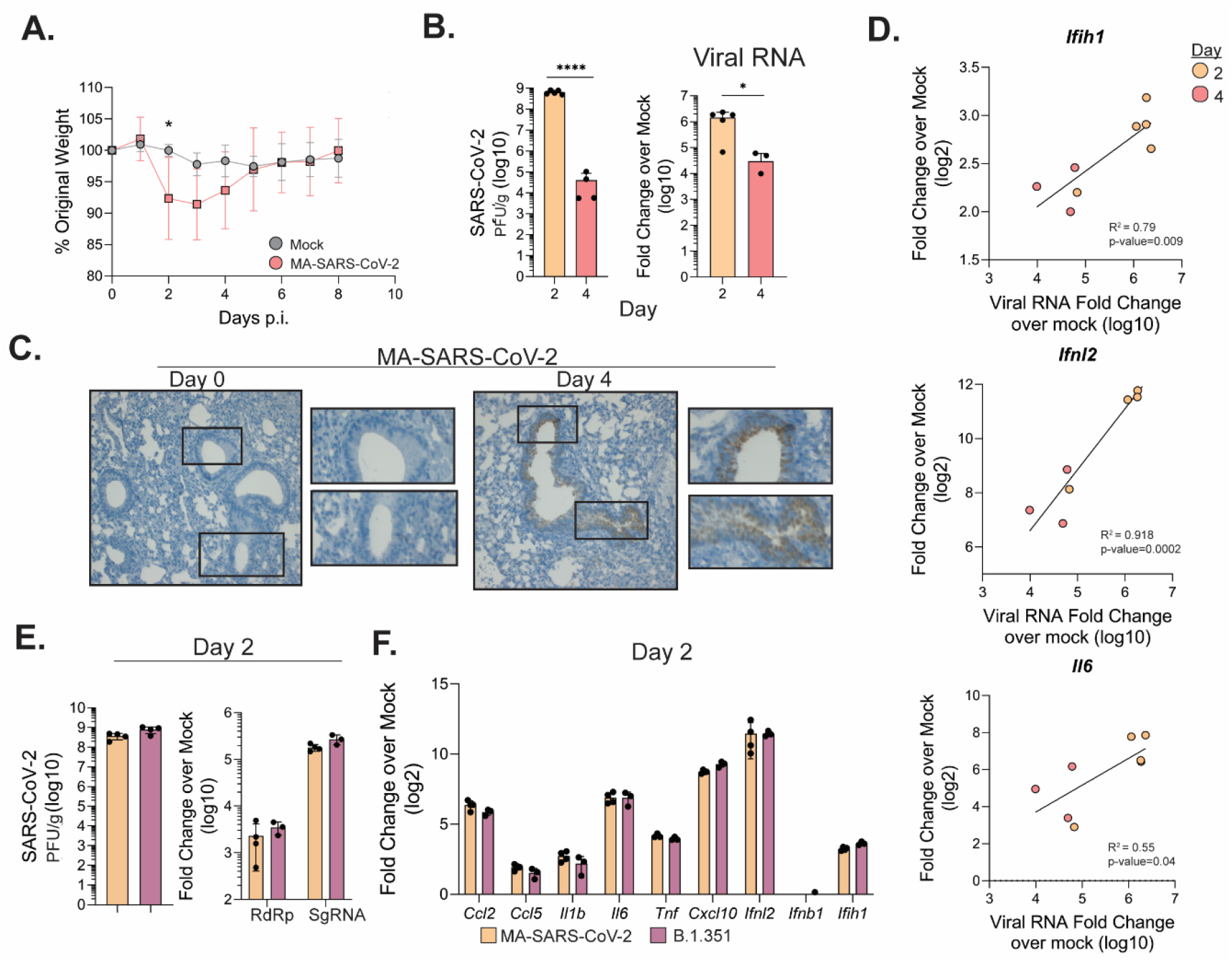
MA-SARS-CoV-2 and B.1.351 replicate in the respiratory tract. C57Bl/6J mice were infected intranasally with 5 x 10^5^ PFU of MA-SARS-CoV-2 or an equal volume of PBS for mock mice. A) Percentage of initial weight for mock and MA-SARS-CoV-2 infected mice over 8 days. B) Quantification of MA-SARS-CoV-2 titers from lung tissue at the indicated day post infection as measured by plaque assay (left) or qRT-PCR (right). CT values are represented as relative fold change over mock (log_10_). C) *In situ* hybridization was performed using a probe for MA-SARS-CoV-2 spike protein RNA. Representative images of lung slices from mock or day 4 p.i. D) Fold-change over mock for the indicated gene was plotted against the corresponding MA-SARS-CoV-2 viral RNA for each sample, and linear regression was used to determine correlation. E) Quantification of viral titers from lung tissue by plaque assay at day 2 p.i. from mice infected with MA-SARS-CoV-2 or human variant B.1.351 (5 x 10^5^ PFU/mouse). On the right, quantification of RNA-dependent RNA-polymerase (RdRp) or subgenomic RNA (SgRNA) fold change over mock. F) Gene expression measured via qRT-PCR for the indicated gene from lungs infected with MA-SARS-CoV-2 or B.1.351 at 2 days p.i. Results are representative of 2 independent experiments with 5 mice per group. Statistical significance was determined using unpaired student’s t-tests or linear regression. * p<0.05, ****p<0.0001.

MA-SARS-CoV-2 contains several mutations, including three within the spike protein, at residues K417N and N501Y, which also appear in the SARS-CoV-2 B.1.351 variant^17^. Next, we evaluated if a natural clinical isolate of B.1.351 could establish infection in mice. We intranasally inoculated C57Bl/6 mice with 5 x 10^5^ PFU of B.1.351 or MA-SARS-CoV-2 and found that B.1.351 replicated to high viral titers within lungs, as measured by plaque assay and qRT-PCR for RNA-dependent RNA-polymerase (RdRp) RNA and subgenomic (Sg) viral RNA (Fig. 1E). The B.1.351 variant induced similar levels of cytokines and ISG expression to MA-SARS-CoV-2 in the lung at day 2 p.i. (Fig. 1F). Combined these data demonstrate that the variant B.1.351 can infect C57Bl/6 mice and replicates similarly to MA-SARS-CoV-2.

### MA-SARS-CoV-2 infection induces hyper-inflammatory monocytes and dysregulated alveolar macrophages

To investigate cellular innate immunity to SARS-CoV-2, we performed single-cell RNA sequencing (scRNA-Seq) on lung homogenates from day 0 or 4 p.i. with MA-SARS-CoV-2 (4 mice per group). We obtained 9,399 day 0 and 10,982 day 4 p.i. cells and unbiased clustering identified 23 distinct groups comprised of T cells, B cells, DCs, epithelial cells (Epi), neutrophils (Neut), natural killer (NK) cells, alveolar macrophages (AM), and monocytes (mono). We further distinguished between inflammatory (Infl), non-classical (NC), and intermediate (Tr) monocytes (Fig. 2A). MA-SARS-CoV-2 induced a decrease in the frequency of epithelial cells and an increase of inflammatory monocytes and DC frequency in the lung (Fig. 2B).

**Figure 2.**
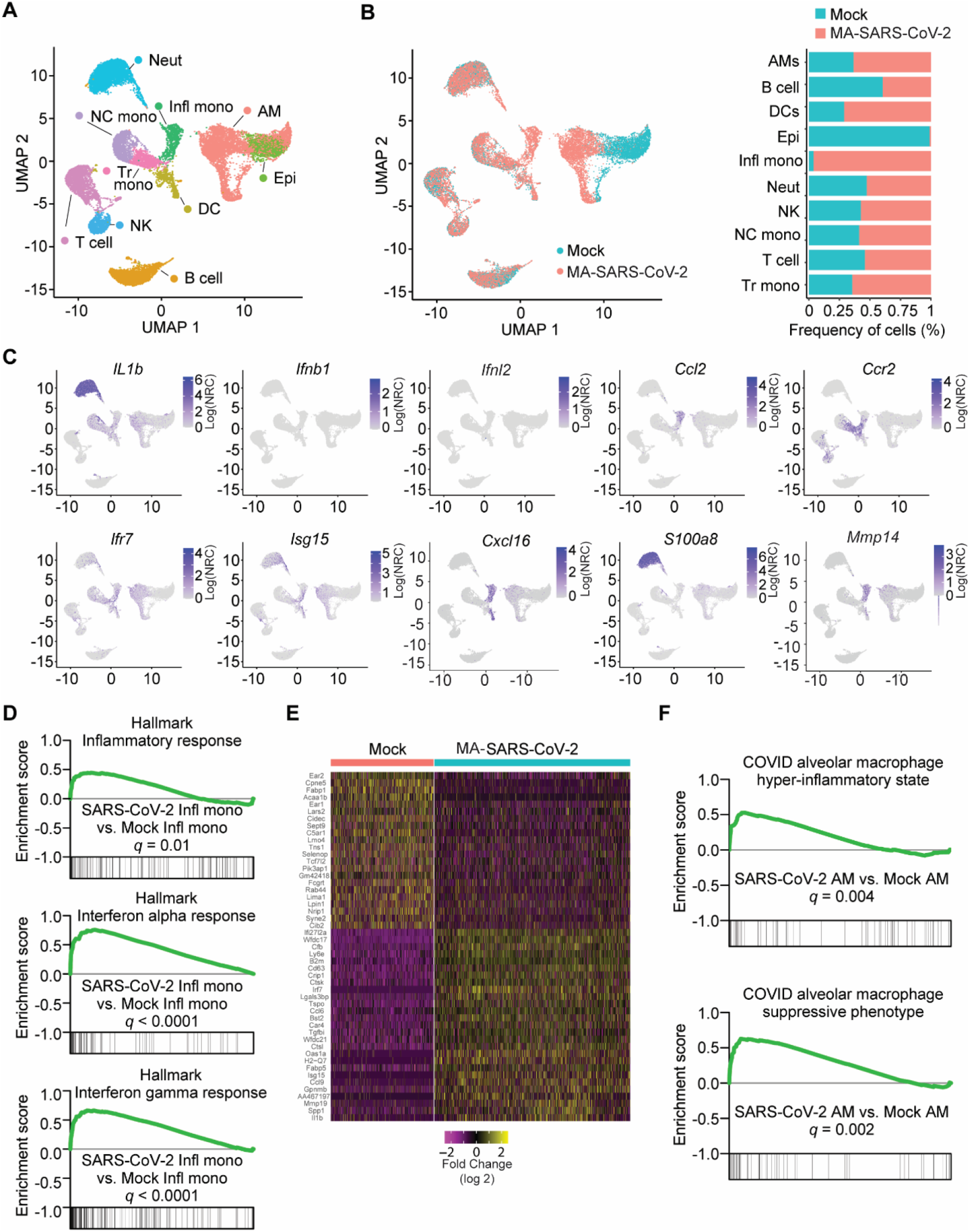
MA-SARS-CoV-2 induces hyper-inflammatory monocytes and macrophages in the lung. C57Bl/6 mice were infected with MA-SARS-CoV-2 and lungs were harvested at day 0 and 4 p.i., processed to a single cell suspension, captured in droplets on a 10X Chromium controller, and analyzed via scRNA-Seq (n=4 per group). A) UMAP plot illustrating the different cellular subsets identified in the lung. B) UMAP distribution of cells from mock or MA-SARS-CoV-2 infected mice. On the right, the frequency of mock vs infected cells that make up each subset defined by UMAP analysis. C) Feature plots displaying the average gene expression of the indicated gene from mock and infected lungs. D) GSEA of inflammatory monocytes using the hallmarks database from MSigDB for the indicated gene set. E) Heatmap analysis of top scoring DEGs in alveolar macrophages from mock or SARS-CoV-2 infected lungs. F) GSEA plots of the indicated gene set from Liao et al. in alveolar macrophages from mock and infected lungs.

We next mapped the expression of genes previously associated with COVID-19 progression to cell subsets in our MA-SARS-CoV-2 model (Fig. 2C). We noted low and sporadic expression of *Ifnb1* or *Ifnl2*, but pronounced ISG expression, *Isg15* and *Irf7*, in alveolar macrophage, monocyte, and DC populations. Inflammatory cytokines were expressed primarily in neutrophils (*Il1b*) or monocytes (*Cxcl16*). Other markers associated with COVID-19, were also localized to neutrophil (*S100a8*) or monocyte (*Mmp14*) populations (Fig. 2C)^18^. While the chemokine *Ccl2* was expressed primarily by inflammatory monocytes, the cognate receptor, *Ccr2*, was more widespread with expression on monocytes, DCs, and NK cells (Fig. 2C). Gene set enrichment analysis (GSEA) of inflammatory monocyte populations identified an enrichment of inflammatory, interferon alpha, and interferon gamma response genes after MA-SARS-CoV-2 infection (Fig. 2D). Thus, MA-SARS-CoV-2 results in a pro-inflammatory response driven by neutrophils and inflammatory monocytes.

Alveolar macrophages showed changes in frequency after MA-SARS-CoV-2 infection (Fig. 2B). To further examine this population, we performed heatmap analysis of top differentially expressed genes (DEGs) between mock and MA-SARS-CoV-2 infected samples. MA-SARS-CoV-2 infection upregulated genes involved in antigen presentation (*B2m, H2-q7*), ISGs (*Ifi2712a, Oas1a*), and inflammatory cytokines (*Ccl9, Ccl6*) (Fig. 2E). We performed GSEA using a gene list enriched in alveolar macrophages from COVID-19 patients and found that alveolar macrophages from MA-SARS-CoV-2 mice had both a hyper-inflammatory and suppressive signature (Fig. 2F)^6^. Together these data demonstrate that alveolar macrophages adopt a dysregulated profile in MA-SARS-CoV-2 infected mice.

### Monocytes and monocyte-derived cells rapidly infiltrate the lung in response to MA-SARS-CoV-2 infection

Next, we investigated the cellular innate immune response to MA-SARS-CoV-2 at days 0, 2, and 4 p.i. Mice were intravitally labelled with CD45 conjugated to phycoerythrin (PE) to allow identification of circulating (CD45+ *in vivo*) and parenchymal (CD45-*in vivo*) cells in the lung. MA-SARS-CoV-2 infection initiated a step-wise increase in total circulating and parenchymal CD45+ cell infiltrate in the lung at days 2 and 4 p.i. (Fig. 3A, See Supplemental Fig. 2 for gating strategy). Granulocyte numbers were elevated in circulation at 2 and 4 days p.i. and infiltrated into the lung by day 2 p.i., with a 100-fold increase in parenchymal neutrophils. (Fig. 3B, Supplemental Fig. 3A). Circulating macrophage numbers were unchanged by infection, however beginning at day 2 p.i., parenchymal macrophage numbers decreased as compared to mock-infected mice (Fig. 3C). This downward trend appeared to be due to a sequential loss of alveolar macrophages at day 2 and 4 p.i. (Siglec-F+ CD11c+), while interstitial macrophages (CD11c-, SiglecF-, Ly6C-) were unaffected (Fig. 3D). At day 4 p.i., we observed a 100-fold increase in cells that expressed both macrophage markers, CD64 and F4/80, and monocyte markers, Ly6C and CD11b, which we designated ‘transitional macrophages’ (Fig. 3D). All macrophages upregulated MHC-I in response to MA-SARS-CoV-2, although the effect was more pronounced in transitional macrophages, which also upregulated CD86 (Supplemental Fig. 3B-C). Together these data identify a shift in the lung macrophage composition, with decreased numbers of alveolar macrophages and increased numbers of activated transitional macrophages during MA-SARS-CoV-2 infection.

**Figure 3.**
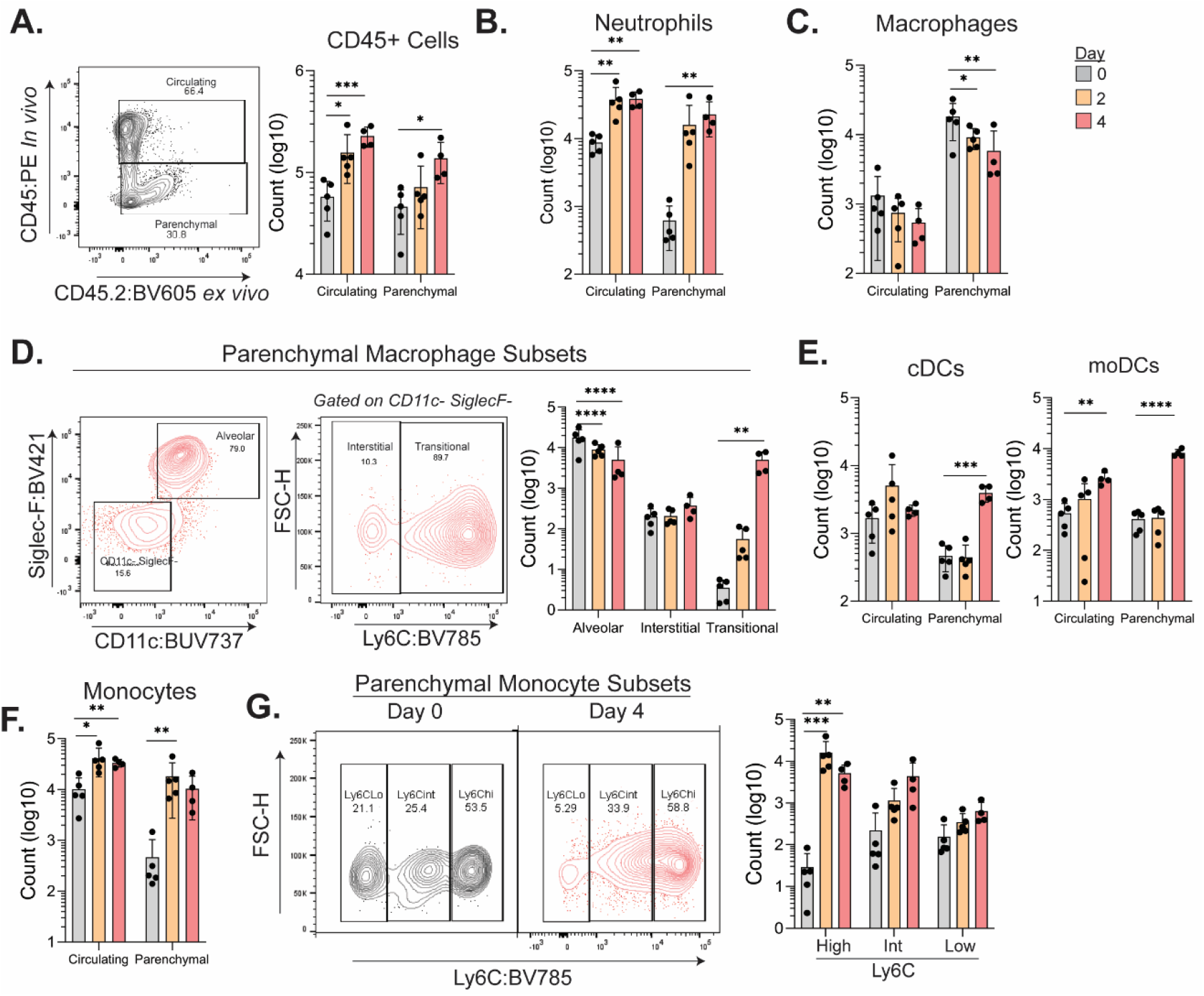
Monocytes and monocyte-derived cells rapidly infiltrate the lung parenchyma in response to MA-SARS-CoV-2 infection. C57Bl/6 mice were infected with MA-SARS-CoV-2 and lung tissue was harvested at 0, 2, and 4 days p.i. and analyzed via flow cytometry. A) 5 minutes prior to euthanization, mice were intravitally labelled with CD45:PE. Representative gating of *in vivo* labelled CD45+ cells used to identify lung circulating (CD45+ *in vivo*) or lung parenchymal (CD45-*in vivo*) cells. The total number of CD45+ *ex vivo* cells is quantified on the right. B) Counts of neutrophils (Lineage-, CD11b+, Ly6G+) over the course of infection. C) Counts of macrophages at day 0, 2, and 4 p.i. (Lineage-, Ly6G-, CD64+, F4/80+). D) Representative flow gating for alveolar (SiglecF+ CD11c+), interstitial (SiglecF- CD11c- Ly6C-), or transitional (SiglecF-, CD11c-, Ly6C+) macrophages from day 4 p.i. Quantified on the right are the counts of each population. E) Quantification of cDCs (lineage-, Ly6G-, CD64-, MHCII+, CD11c+, CD26+) or moDCs (lineage, Ly6G-, MHC-I I+, CD11b+, CD11c+) counts at the indicated timepoint. F) Total monocyte (lineage-, Ly6G-, MHC-II-, CD11c-, CD64 low) counts for circulating or lung parenchymal cells at day 0, 2, and 4 p.i. G) Representative gating strategy to identify different monocyte subsets (Ly6C High, Ly6C Intermediate (Int), Ly6C low), and the quantification on the right. Results are representative of two independent experiments with 5 mice per group. Statistical significance was determined using unpaired one or two-way ANOVA. *p<0.05, **p<0.01, ***p<0.001, ****p<0.0001.

We next examined the role of dendritic cells during MA-SARS-CoV-2 infection. Plasmacytoid dendritic cell (pDC) numbers did not change in the circulation or parenchyma at 2-4 days p.i. (Supplemental Fig. 3D). Conventional dendritic cell (cDC) populations remained steady in circulation, but increased 10-fold in the lung parenchyma at day 4 p.i. that was primarily due to an increase in cDC Type 2 cells (cDC2s) (Fig. 3E, Supplemental Fig. 3E). Lung parenchymal cDCs increased expression of MHC-I and CD86 in response to MA-SARS-CoV-2 at day 4 p.i (Supplemental Fig. 3F). moDCs had slightly increased numbers in circulation and a 10-fold increase in lung parenchymal populations at day 4 p.i. (Fig. 3E). Parenchymal moDCs also upregulated expression of MHC-I and CD86 at day 4 p.i. as compared to uninfected controls (Supplemental Fig. 3G). Thus, cDCs and moDCs undergo expansion and activation in response to MA-SARS-CoV-2 infection in the lung.

Investigation of monocyte dynamics during MA-SARS-CoV-2 infection observed a 5-fold increase in circulating monocytes and a 20-fold increase in lung parenchymal monocytes by day 2 p.i. that remained high through day 4 p.i. (Fig. 3F). Ly6C high monocytes drove monocytic infiltration to the lung, as their numbers increased 100-fold at days 2 and 4 p.i., but the numbers of Ly6C low monocytes remained constant (Fig. 3G). All lung parenchymal monocytes showed increased expression of MHC-I at day 4 p.i. (Supplemental Fig. 3H). Ly6C high monocytes had particularly elevated expression of CD86 (2000% increase) at day 4 p.i. as compared to mock (Supplemental Fig. 3I). Analysis of splenic immunity found that neutrophils had significantly increased numbers at day 4 p.i. (Supplementary Fig. 4A). Dendritic cells showed increased expression of MHC-I at 4 days p.i. (Supplementary Fig. 4B). Together these data find that MA-SARS-CoV-2 infection prompts systemic immune activation and a lung parenchymal immune response dominated by the infiltration of activated monocytes and monocyte-derived cells.

### Expansion of monocyte-derived cells in the lung parenchyma during MA-SARS-CoV-2 infection is CCR2-dependent

Classical Ly6C high monocytes migrate to the lung parenchyma in a CCR2-dependent manner and can differentiate into interstitial macrophages or moDCs^11,19^. We next evaluated the contribution of CCR2 signaling in recruiting monocytes to the lung during MA-SARS-CoV-2 infection. Flow cytometry analysis of lungs at day 4 p.i. found similar circulating monocyte numbers, but a 2-fold drop in the number of lung parenchymal infiltrating monocytes in *Ccr2^-/-^* mice as compared to WT (Fig. 4A). This decrease appeared to be driven by a 4-fold drop in Ly6C high and Ly6C intermediate monocytes in the lungs of *Ccr2^-/-^* mice (Fig. 4B). Circulating moDC numbers were unaltered by the absence of CCR2, but lung parenchymal moDCs dropped 10-fold in *Ccr2^-/-^* mice at day 4 p.i. (Fig. 4C). cDC numbers in circulation were similar between WT and *Ccr2^-/-^* mice at day 4 p.i., but lung parenchymal cDC numbers at day 4 p.i. were 5-fold lower than WT (Fig. 4D). This was due to a specific loss in cDC2s (Fig. 4D). All monocyte subsets showed decreased expression of CD86 (2-fold decrease) at day 4 p.i. in *Ccr2^-/-^* mice, while MHC-I levels were decreased only in the Ly6C intermediate subset (Fig. 4E). Expression of antigen presentation markers on moDCs was unchanged by CCR2 (Supplemental Fig. 5A). Lung-infiltrating cDC2s from *Ccr2^-/-^* mice had lower expression of MHC-I, but not CD86 as compared to WT cells at 4 days p.i. (Supplemental Fig. 5B). Together these data find that CCR2 signaling promotes the infiltration of activated Ly6C high and intermediate monocytes, moDCs, and cDC2s into the lung parenchyma during MA-SARS-CoV-2 infection.

**Figure 4.**
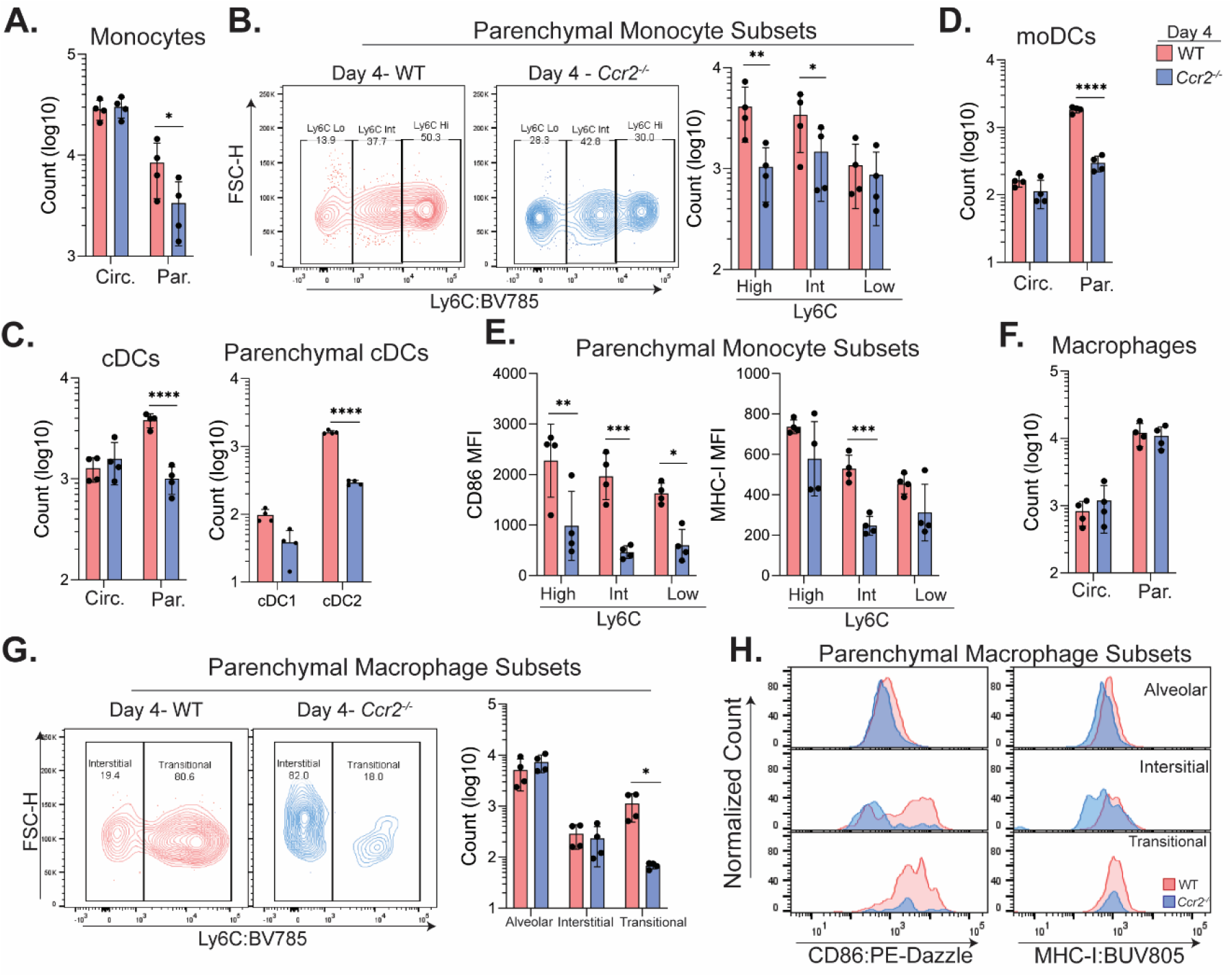
Expansion of monocyte-derived cells in the lung during MA-SARS-Cov-2 is CCR2 dependent. C57Bl/6 and *Ccr2^-/-^* mice were infected with MA-SARS-CoV-2 and lung tissue was harvested at 0 and 4 days p.i. and analyzed via flow cytometry. Circulating (Circ.) vs parenchymal (Par.) cells were distinguished as described in Fig. 2. A) Number of total monocytes in the lung circulation or parenchyma B) Representative gating identifying Ly6C High, Intermediate (Int), or Low monocytes from WT and *Ccr2^-/-^* lung parenchyma. Counts for each subset are quantified to the right. C) Quantification of moDCs at day 4 p.i. D) Total numbers of cDCs and parenchymal cDC subsets (right) E) MFI for CD86 (left) and MHC-I (right) expression on monocyte subsets from the lung parenchyma. F) Quantification of the total macrophage numbers at day 4 p.i. G) Representative flow plots illustrating interstitial or transitional macrophage populations from WT and *Ccr2^-/-^* lung infiltrating cells. Counts of macrophage subsets are quantified on the right. H) Representative histograms of the MFI for CD86 (left) and MHC-I (right) for each macrophage subset 4 days p.i. from WT or *Ccr2^-/-^* lung infiltrating cells. Results are representative of two independent experiments with 5 mice per group. Statistical significance was determined using unpaired one or two-way ANOVA. *p<0.05, **p<0.01, ***p<0.001, ****p<0.0001.

We investigated if CCR2 low/negative innate immune cells (Fig. 3C) were impacted by secondary effects of CCR2. The total number of macrophages in circulation or in the lung parenchyma was not affected by CCR2 at 4 p.i. with MA-SARS-CoV-2 (Fig. 4F). Alveolar and interstitial macrophages were not impacted by CCR2 signaling. However, *Ccr2^-/-^* mice had a 10-fold drop in transitional macrophage numbers as compared to WT mice (Fig. 4G). Expression of CD86 and MHC-I was also decreased on transitional macrophages from *Ccr2^-/-^* mice at day 4 p.i. Despite similar numbers in WT and *Ccr2^-/-^* mice, interstitial macrophages failed to upregulate expression of both MHC-I and CD86 in *Ccr2^-/-^* mice at day 4 p.i. Expression of CD86 MFI on alveolar macrophages from *Ccr2^-/-^* mice was modestly decreased compared to WT (Fig. 4H). Thus, activation of macrophages and expansion of transitional macrophages is CCR2 dependent during MA-SARS-CoV-2 infection.

Lung parenchymal granulocyte numbers were not significantly altered between WT and *Ccr2^-/-^* mice at day 4 p.i. (Supplemental Fig. 5C). However, there was a modest increase in the number of granulocytes in circulation and the spleen from *Ccr2^-/-^* mice as compared to WT mice at day 4 p.i. (Supplemental Fig. 5C-D). Total splenic macrophage numbers were also higher in the absence of CCR2 at day 4 p.i. (Supplemental Fig. 5D). Monocyte, moDC, and pDC populations in the spleen were unaffected by CCR2, however numbers of cDCs in the spleen were increased in *Ccr2^-/-^* mice at day 4 p.i. as compared to WT (Supplemental Fig. 5D). These data identify a role for CCR2 in promoting the infiltration of activated, Ly6C high monocytes and monocyte derived cells to the lung during MA-SARS-CoV-2 infection.

### CCR2 signaling restricts MA-SARS-CoV-2 in the lung

To determine if CCR2 was protective against MA-SARS-CoV-2, we infected WT or *Ccr2^-/-^* mice with MA-SARS-CoV-2 and assessed viral burden at day 4 p.i. *Ccr2^-/-^* mice had 10-fold higher viral burden in lung tissue as measured by plaque assay or qRT-PCR (Fig. 5A-B). At day 4 p.i., *Ifnl2*, cytokines (*Il6*), and chemokines (*Cxcl10, Ccl2*) were elevated in *Ccr2^-/-^* as compared to WT lungs (Fig. 5C). *In situ* hybridization found that *Ccr2^-/-^* lungs had more robust detection of viral RNA than WT mice. Additionally, while viral RNA was localized to cells lining the airway spaces in WT mice, *Ccr2^-/-^* lungs had infiltration of viral RNA further into the interstitial and parenchymal spaces (Fig. 5D). Together, these data find that CCR2 mediated signaling and immune cell recruitment restrict inflammation, viral burden, and viral dissemination in the lung during MA-SARS-CoV-2 infection.

**Figure 5.**
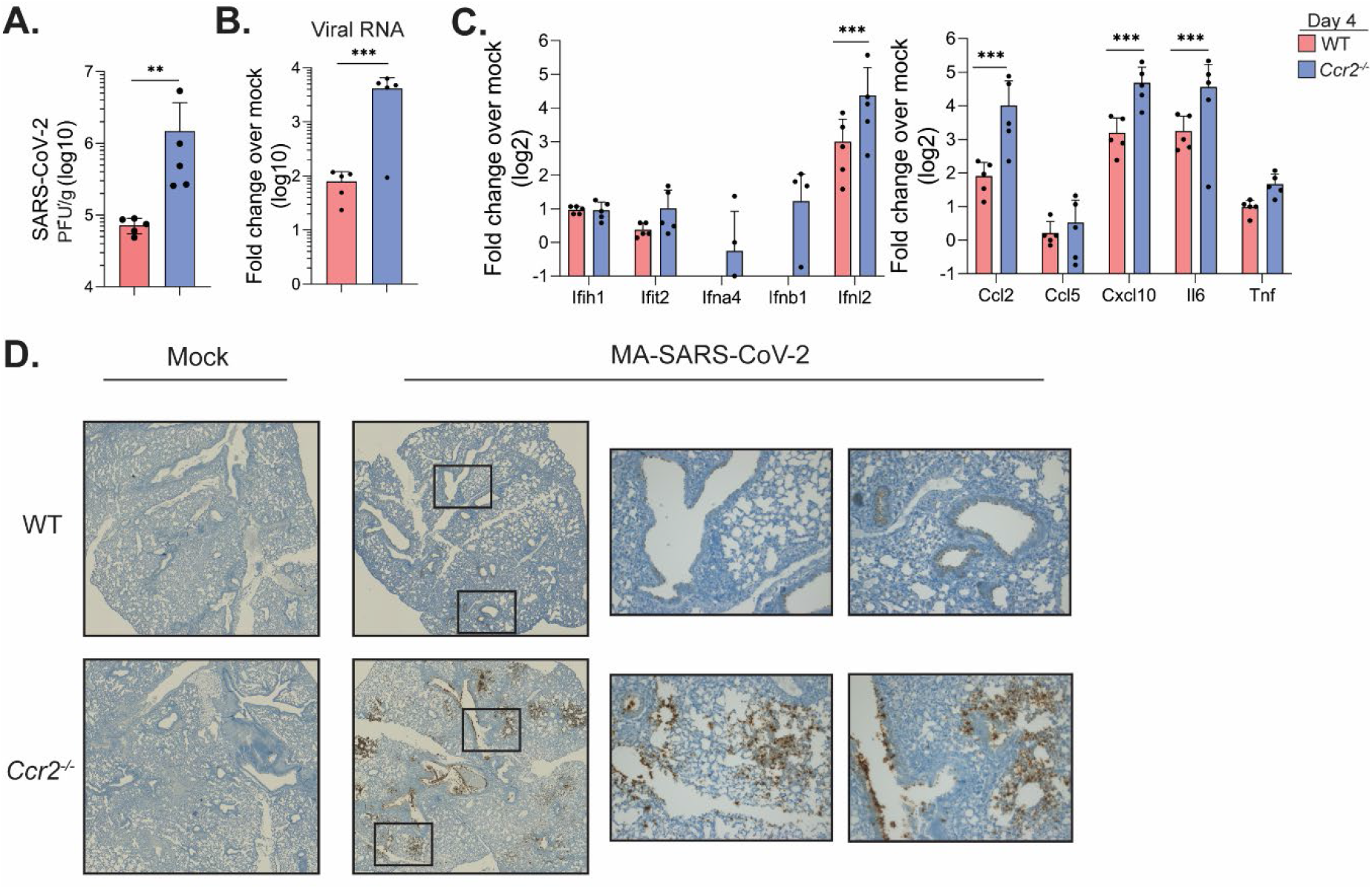
CCR2 restricts MA-SARS-CoV-2 viral burden and inflammatory cytokines in the lung. C57Bl/6 or *Ccr2^-/-^* mice were infected with MA-SARS-CoV-2 and lung tissue was collected at day 4 p.i. A) Infectious virus at day 4 p.i. as quantified via plaque assay. B) qRT-PCR for the SARS-CoV-2 RdRp. C) qRT-PCR was performed to probe for the indicated IFN signaling (left) or inflammatory (right) transcripts. D) Representative images of *in situ* hybridization to visualize MA-SARS-CoV-2 RNA in lung tissue slices from 0 and 4 days p.i. in both WT and *Ccr2^-/-^* mice. Results are representative of two independent experiments with 5 mice per group. Statistical significance was determined using unpaired student’s t-test or one-way ANOVA. *p<0.05, **p<0.01, ***p<0.001, ****p<0.0001.

## Discussion

SARS-CoV-2 infection in humans has identified a robust and reproducible correlation between inflammatory cytokine levels and disease severity during COVID-19^6^. In accordance, we found that SARS-CoV-2 infected mice had a pro-inflammatory cytokine profile in the lung containing pyrogens (*Il6, Tnf*), chemoattractants for monocytes (*Ccl2*) and T cells (*Cxcl10*), ISGs (*Irf7* and *Isg15*), alarmins (*S100a8*), and matrix metalloproteinases (*Mmp14*)^6,20–23^. Interestingly, inflammatory gene expression in the lung was MA-SARS-CoV-2 viral load dependent, as most cytokine transcripts surveyed by qPCR positively correlated with viral RNA. Similar phenomenon have been noted in human subjects, with one study describing an association between SARS-CoV-2 viral burden, IL-6 levels, and increased risk of death^24^. Similar to studies of post-mortem lung tissue or BALs from patients suffering from COVID-19, we observed a significant increase in the numbers of *S100a8+* granulocytes in the lung parenchyma^4,6,8^. S100a8/9 and TLR4 signaling mediate the emergence of a dysregulated neutrophil population that promotes SARS-CoV-2 disease^18^. Neutrophil and inflammatory cytokine levels were higher in *Ccr2^-/-^* mice, likely driven by increased viral burden in *Ccr2^-/-^* lungs. Thus, MA-SARS-CoV-2 viral burden is likely directly driving cytokine expression and neutrophilic infiltration into the lung.

CCR2 expression is a defining feature of inflammatory Ly6C high blood monocytes^9^. In various models of respiratory disease including influenza, *Mycobacterium tuberculosis*, and allergic inflammation, deletion of *Ccr2* negates the ability of monocytes to enter the lung parenchyma^11,25,26^. Similarly, we found that CCR2 had no effect on circulating monocyte numbers but specifically promoted infiltration of activated monocytes into the lung parenchyma during SARS-CoV-2. Monocyte-derived interstitial macrophages and moDCs also depended on CCR2 for expansion in the lung parenchyma^27^. In contrast, alveolar macrophages, which are self-renewing and primarily fetal derived, were unaffected by the absence of CCR2^28–30^. Previous studies in a model of sterile lung inflammation demonstrated that DC precursors required CCR2 for entry to the lung, and accordingly in MA-SARS-CoV-2 infection, cDC2 lung parenchymal populations were also decreased in the absence of CCR2^31^. These data suggest that CCR2 plays an essential and specific role in promoting the recruitment and differentiation of monocytes into transitional macrophage and moDC populations during MA-SARS-CoV-2 infection.

In contrast to other respiratory infections such as influenza, in which CCR2 promotes mortality and inflammation, our study found that CCR2 restricted viral burden and weight loss during MA-SARS-CoV-2 infection^11^. CCR2 promoted the infiltration of monocyte-derived cells into the lung parenchyma, thus suggesting an essential role for Ly6C high monocytes, moDCs, and transitional macrophages in controlling SARS-CoV-2 infection. Macrophages have been previously shown to control viruses through the secretion of iNOS and the phagocytosis of viral particles, while moDCs play a key role in Type I IFN release and priming the T cell response^13,32^. Further studies are needed to delineate the precise mechanisms these cell subsets use to control SARS-CoV-2. Current evidence in the field of SARS-CoV-2 research suggests that a large portion of the pathology of COVID-19 is due to a monocyte-driven cytokine storm^33^. Here we identify data that suggests monocyte-derived cells in the lung are crucial for limiting viral burden and cytokine production during the early stages of MA-SARS-CoV-2 infection. Most studies to date of the immune response in humans have focused on late-timepoints of infection, therefore monocytes could be protective early and pathological later during infection resolution. The consideration of the role of monocyte-derived cells in restricting SARS-CoV-2 infection in the lung and priming adaptive immune responses will be essential for the design of future therapies and vaccines.

## METHODS

### Viruses and cells

VeroE6 cells were obtained from ATCC (clone E6, ATCC, #CRL-1586) and cultured in complete DMEM medium consisting of 1x DMEM (VWR, #45000-304), 10% FBS, 25mM HEPES Buffer (Corning Cellgro), 2mM L-glutamine, 1mM sodium pyruvate, 1x Non-essential Amino Acids, and 1x antibiotics. VeroE6-TMPRSS2-hACE2 cells were kindly provided by Dr. Barney Graham (Vaccine Research Center, NIH, Bethesda, MD). The generation of the MA-SARS-CoV-2 virus will be described in a future publication. In brief, MA-SARS-CoV-2 virus was generated by engineering four coding mutations (NSP6 L37F, NSP10 P87S, S N501Y, and N D128Y) into the backbone of icSARS-CoV-2 (WA/1 backbone). This virus was then passaged 20 times in Balb/c mice followed by deep sequencing which identified 3 additional acquired mutations (S T417N, S H655Y, and E E8V). This virus, termed MA-SARS-CoV-2 was passaged once in VeroE6 cells to generate a working stock. The B.1.351 variant was provided by Dr. Andy Pekosz (John Hopkins University, Baltimore, MD) and grown on VeroE6-TMPRSS2 cells and viral titers were determined by plaque assay on VeroE6 cells (ATCC). The generation of TMPRSS2-hACE2 cells were previously described^34^. VERO E6 cells (ATCC CRL-1586) were transfected with pCAGGS plasmid in which chicken actin gene promoter drives the expression of an open reading frame comprising Puromycin N-acetyl transferase, GSG linker, 2A self-cleaving peptide of thosea asigna virus (T2A), human transmembrane serine protease 2 (TMPRSS2). Two days post-transfection, cells were trypsinzed and transferred to a 100 mm dish containing complete DMEM medium (1x DMEM, Thermo Fisher, # 11965118, 10% FBS, 1x penicillin/streptomycin) supplemented with puromycin (Thermo Fisher, #A1113803) at a final concentration of 10 μg/ml. Approximately ten days later, individual colonies of cells were isolated using cloning cylinders (Sigma) and expanded in medium containing puromycin. Clonal cell lines were screened for expression of TMPRSS2 by flow cytometry. VeroE6-TMPRSS2 cells and VeroE6-TMPRSS2-hACE2 cells were cultured in complete DMEM in the presence of Gibco Puromycin 10mg/mL (#A11138-03). VeroE6 cells were cultured in complete DMEM medium consisting of 1x DMEM (Corning Cellgro), 10% FBS, 25 mM HEPES Buffer (Corning Cellgro), 2 mM L-glutamine, 1mM sodium pyruvate, 1x Non-essential Amino Acids, and 1x antibiotics.

### Infection of mice with MA-SARS-CoV-2

C57BL/6J and *Ccr2^-/-^* mice were purchased from Jackson Laboratories or bred in-house at the Yerkes National Primate Research Center rodent facility at Emory University. All mice used in these experiments were females between 8-12 weeks of age. Stock MA-SARS-CoV-2 or B.1.351 virus was diluted in PBS to a working concentration of 1 x 10^7^ PFU/mL. Mice were anesthetized with isoflurane and infected intranasally with virus (50 uL, 5 x 10^5^ PFU/mouse) in an ABSL-3 facility. Mice were monitored daily for weight loss. All experiments adhered to the guidelines approved by the Emory University Institutional Animal Care and Committee.

### Quantification of infectious virus

At the indicated day post infection, mice were euthanized via isoflurane overdose and lung tissue was collected in Omni-Bead ruptor tubes filled with 1% FBS-HBSS. Tissue was homogenized in an Omni Bead Ruptor 24 (5.15 ms, 15 seconds). To perform plaque assays, 10-fold dilutions of viral supernatant in serum free DMEM (VWR, #45000-304) were overlaid on VeroE6-TMPRSS2-hACE2 cells monolayers and adsorbed for 1 hour at 37°C. After adsorption, 0.5% immunodiffusion Agarose in 2X DMEM supplemented with 5% FBS (Atlanta Biologics) and 1X sodium bicarbonate was overlaid, and cultures were incubated for 48 hours at 37°C. Agarose plugs were removed, cells fixed with 4% PBS-buffered paraformaldehyde for 15 minutes at room temperature and plaques were visualized using crystal violet staining (20% methanol in ddH_2_O).

### Quantitative reverse-transcription PCR of lung tissues

At the indicated day post infection, mice were euthanized with isoflurane overdose and one lobe of lung tissue was collected in an Omni Bead ruptor tube filled with Tri Reagent (Zymo, #R2050-1-200). Tissue was homogenized using an Omni Bead Ruptor 24 (5.15 ms, 15 seconds), then centrifuged to remove debris. RNA was extracted using a Direct-zol RNA MiniPrep Kit (Zymo, # R2051), then converted to cDNA using a High-capacity Reverse Transcriptase cDNA Kit (Thermo, #4368813). RNA levels were quantified using the IDT Prime Time Gene Expression Master Mix, and Taqman gene expression Primer/Probe sets (IDT). All qPCR was performed in 384-well plates and run on a QuantStudio5 qPCR system. SARS-CoV-2 RNA-dependent RNA polymerase levels were measured as previously described^5^. SARS-CoV-2 E protein subgenomic RNA (sgRNA) was measured using the E_Sarbeco set from IDT: F1 forward primer, 100 nmol (#10006889), R2 reverse primer, 100 nmol (#10006891), P1 probe, 50 nmol (#10006893). The following Taqman Primer/Probe sets (ThermoFisher) were used in this study: Gapdh (Mm99999915_g1), Ifnl2 (Mm04204155_gH), Ifit2 (Mm00492606_m1), Ifih1(Mm00459183_m1), Ifnb1 (Mm00439552_s1), Ifna4 (Mm00833969_s1), Ccl2 (Mm00441242_m1), Ccl5 (Mm01302427_m1), Cxcl10 (Mm99999072_m1), Tnf (Mm00443258_m1), Il1b (Mm00434228_m1), Il6 (Mm00446190_m1).

### Processing of mouse tissues to single cell suspensions

At the indicated day post infection, mice were anesthetized using isoflurane and injected retro-orbitally with CD45:PE (100 ul per mouse, diluted 1:20 in PBS). Mice were allowed to recover for 5 minutes, then euthanized via isoflurane overdose. One lobe of lung tissue and spleens were collected from each mouse and placed in 1% FBS-HBSS. Spleens were mechanically homogenized on a 70 uM cell strainer, and the cell suspension was collected in 10% FBS-RPMI. Splenocyte suspension was spun down (1250 rpm, 5 min, 4C) and lysed in ACK Lysis buffer (Lonza) for 5 minutes on ice. Splenocytes were washed with 10%FBS-RPMI then kept on ice until ready for downstream applications. Lungs were mechanically disrupted in 6-well plates, then digested for 30 minutes at 37C in a solution of DNaseI (2000U/mL, Sigma D4527-500KU)) and collagenase (5mg/mL, (Sigma, #11088882001) in HBSS. Digestion was stopped with 10% FBS-RPMI, and lungs were pushed through a 70 uM filter to obtain a single cell suspension. Cells were resuspended in 30% Percoll-PBS and centrifuged at 2000 rpm for 20 minutes. The top layer of cell debris was removed and the cell-pellet at the bottom was lysed with ACK lysis buffer for 5 minutes on ice. Cells were washed, and resuspended in 10%FBS-RPMI and kept on ice until ready for staining.

### Flow cytometry analysis

Single-cell suspensions were spun down, and resuspended in anti-CD16/32 (Tonbo, Clone 2.4G2) blocking solution for 20 minutes at 4°C. Cell suspensions were spun down, and stained with Live/Dead Ghost Dye stain (Tonbo Biosciences) for 20 minutes at 4°C. Cells were washed, and resuspended in the indicated surface stain in FACS buffer for 20 minutes at 4°C. After staining, cells were washed and fixed in 2% PFA-PBS for 20 minutes at room temperature. Precision count beads (Biolegend) were added to samples to obtain counts. Samples were run on a BD FACS Symphony A5. The following antibodies were used in this study: CD45:PE (Biolegend, Clone: 30-F11), CD45.2:BV605 (Biolegend, Clone: 104), CD11b:BUV395 BD Biosciences, 440c), I-A/I-E:AF700 (Biolegend, M5/114.15.2), CD11c:BUV737 (Biolegend, N418), CD26: PE-Cy7 (Biolegend, H194-112), CD172a:BV510 (Biolegend, P84), XCR1:AF647 (Biolegend, Zet), CD64: PerCpCy5.5 (Biolegend, X54-5/7.1), F4/80:FITC (Biolegend, BM8), H2kb:BUV805 (BD Biosciences, AF6-88.5), CD86:PE-Dazzle594 (Biolegend, GL1), Live/Dead Ghost Dye:R780 (Tonbo), CD3:APC-Cy7 (Biolegend, 145-2C11), CD19:APC-Cy7 (BD Biosciences, 1D3), NK1.1 (Biolegend, PK136), Ly6C:BV785 (Biolegend, Hk1.4), Ly6G:BV650 (Biolegend, 1A8), SiglecF:BV421 (BD Biosciences, E50-2440).

### *In situ* hybridization of lung tissues

One lobe of the lung was harvested from mice at 0 or 4 days p.i., and fixed with 4% PFA-PBS for a minimum of 3 days. Formalin-fixed paraffin-embedded lung tissue were deparaffinized through sequential washes twice each in xylene and 100% ethanol for 5 min. Tissue were then pretreated with RNAscope Hydrogen Peroxide for 10 min at RT, then RNAscope Target Retrieval for 5 min at 95-100 °C, followed by RNAscope Protease Plus for 30 min at 40 °C. RNA-ISH was performed using a probe against the S gene of SARS-CoV-2 (V-nCoV-2019-S, ACD) using the RNAscope 2.5 HD Assay-BROWN as per the manufacturer’s instructions. Slides were covered slipped with ProLong Gold Antifade Mountant (Thermo Fisher). Images were acquired using a Zeiss AxioImager Z2 system with Zeiss software.

### Single-cell RNA-Seq analysis

Lungs from mice at 0 or 4 days p.i., were processed to single cells suspensions as described above. Single cell suspensions were washed 4 times with PBS, and passed through a 70 uM filter. Cell suspensions were counted, and captured in droplets using the Chromium NextGEM Single Cell 5’ Library & Gel Bead kits on a 10x Chromium controller in a BSL-3 biosafety cabinet. Amplification of cDNA and library preparation was performed according to the manufacturer’s instructions. Gene expression libraries were sequenced as paired-end 26×91 reads on an Illumina NovaSeq6000 targeting a depth of 50,000 reads per cell in the Yerkes Genomics Core Laboratory (http://www.yerkes.emory.edu/nhp_genomics_core/). Analysis was conducted using R (v4) and Seurat (v4). Cell Ranger (v6) was used for demultiplexing, aligning barcodes, mapping to the genome (mm10) and quantifying UMIs. Filtered Cell Ranger matrices were processed with Read10x function in Seurat for preprocessing and cluster analysis. Data was filtered to remove cells with less than 200 genes, abnormally high gene counts (feature counts >5000) and greater than 5% mitochondrial genes. After qualify control, there were 9,399 mock cells and 10,982 CoV2 cells. Principal component analysis (PCA) and dimensional reduction was conducted on Log-normalized and scaled gene expression data. Clustering was conducted using FindNeighbors and FindClusters functions, with resolution parameters between 0.5-1.4. Overall, 23 clusters were identified and FindAllMarkers function utilized to identify DEGs, from which marker genes for cluster cell annotation was conducted. After annotating cells, DEGs were determined based on sub-clusters or experimental group. Gene set enrichment analysis (GSEA) was conducted using ranked gene list produced with Seurat FindMarkers function (comparing CoV2 samples with mock samples) and Genelist were obtained from: MsigDB (hallmarks) and https://www.nature.com/articles/s41422-020-00455-9#MOESM1 (macrophage suppressive and hyperinflammatory). GSEA was conducted using Broad (4.0.3) and plotted in R.

## Data availability

Single cell RNA sequencing data will be publicly accessible through Gene Expression Omnibus following acceptance of this article.

## Acknowledgements

This work was supported in part by grants (P51 OD011132 and R56 AI147623 to Emory University) from the National Institute of Allergy and Infectious Diseases, National Institutes of Health, by intramural funding from the National Institute of Allergy and Infectious Diseases, the Emory Executive Vice President for Health Affairs Synergy Fund award, the Pediatric Research Alliance Center for Childhood Infections and Vaccines and Children’s Healthcare of Atlanta, the Emory-UGA Center of Excellence for Influenza Research and Surveillance, and Woodruff Health Sciences Center 2020 COVID-19 CURE Award. J.T. is supported by T32 AI074492. We thank Andy Pekosz for the B.1.351 variant. Next-generation sequencing services were provided by the Yerkes NHP Genomics Core, which is supported in part by NIH P51 OD 011132, and the data were acquired on a NovaSeq 6000 funded by NIH S10 OD 026799. We also thank Eli Boritz and Danny Douek for sequencing and analysis of the B.1.351 variant (NIAID/NIH, Atlanta, GA).

## Author Contributions

A.V. and M.S.S. contributed to the acquisition, analysis, and interpretation of the data, as well as the conception and design of the work, and writing the manuscript. J.T. and J.K. contributed to the analysis and interpretation of the data, and writing the manuscript. A.S. contributed to the acquisition, analysis, and interpretation of the data, and writing the manuscript. M.D.G, K.F., A.C, A.P, A.D-C, and F.J. contributed to the acquisition and analysis of the data. R.K. contributed to the analysis, interpretation of the data, and conception and design of the work. S.E.B., D.A.C, K.P., A.G., P.S. and V.D.M. contributed to the interpretation of the data and conception and design of the work.

## Supplemental Figures

**Supplemental Figure 1.**
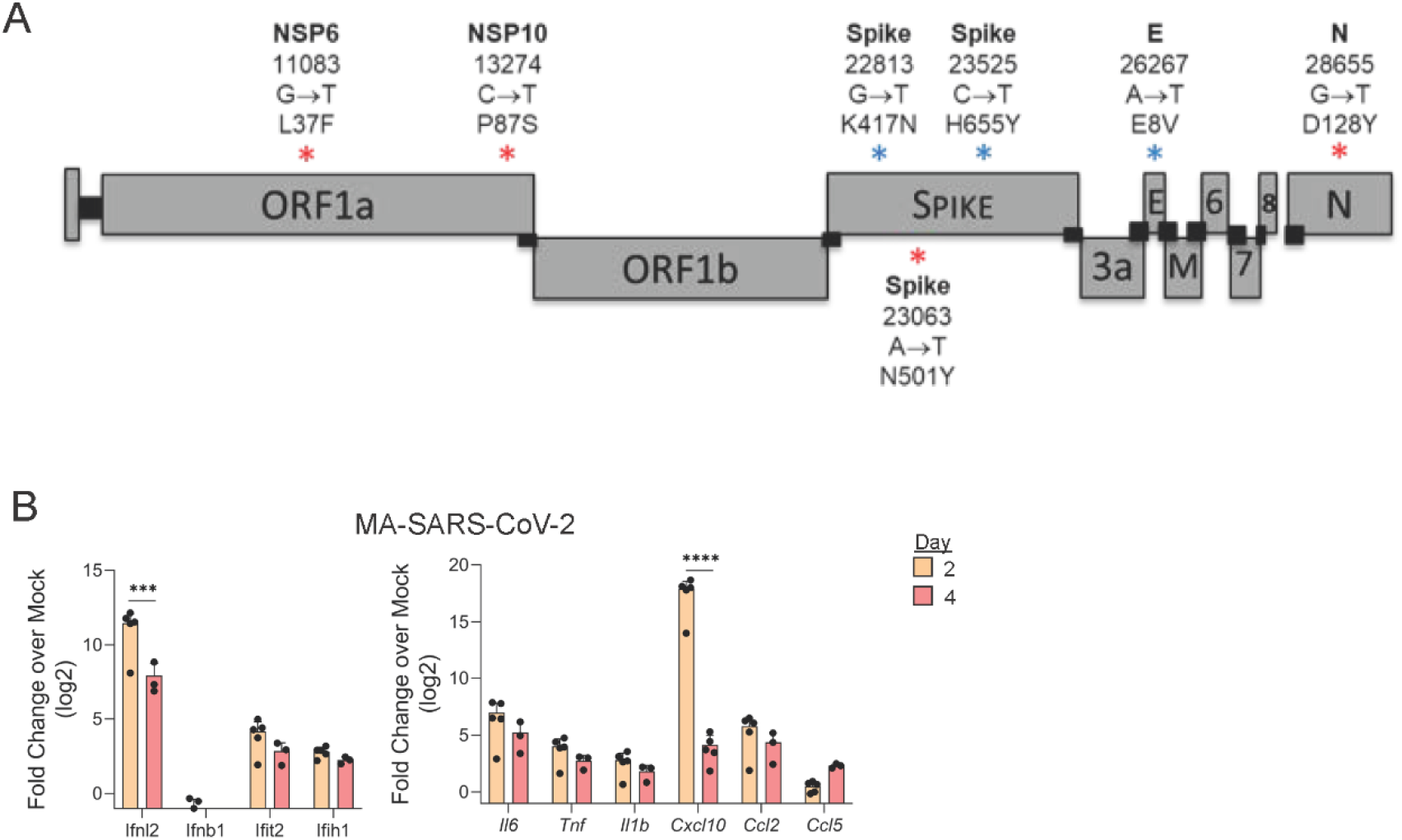
MA-SARS-CoV-2 induces an inflammatory response in the lung. A) Schematic of MA-SARS-CoV-2. Mutations with a red asterisk were engineered into icSARS-CoV-2 followed by passaging the virus in Balb/c mice. Following 20 passages, the virus was sequenced and mutations with a blue asterisk were acquired in addition to the engineered mutations during the passage of MA-SARS-CoV-2. B) Mice were treated with PBS or infected with MA-SARS-CoV-2 via the intranasal route and lung tissue was harvested from 0, 2, and 4 days p.i. Fold change over mock for the indicated genes at day 2 or 4 p.i. with MA-SARS-CoV-2. Results are representative of 2 independent experiments. Statistical significance was determined using unpaired two-way ANOVA. ***p<0.001, ****p<0.0001.

**Supplemental Figure 2.**
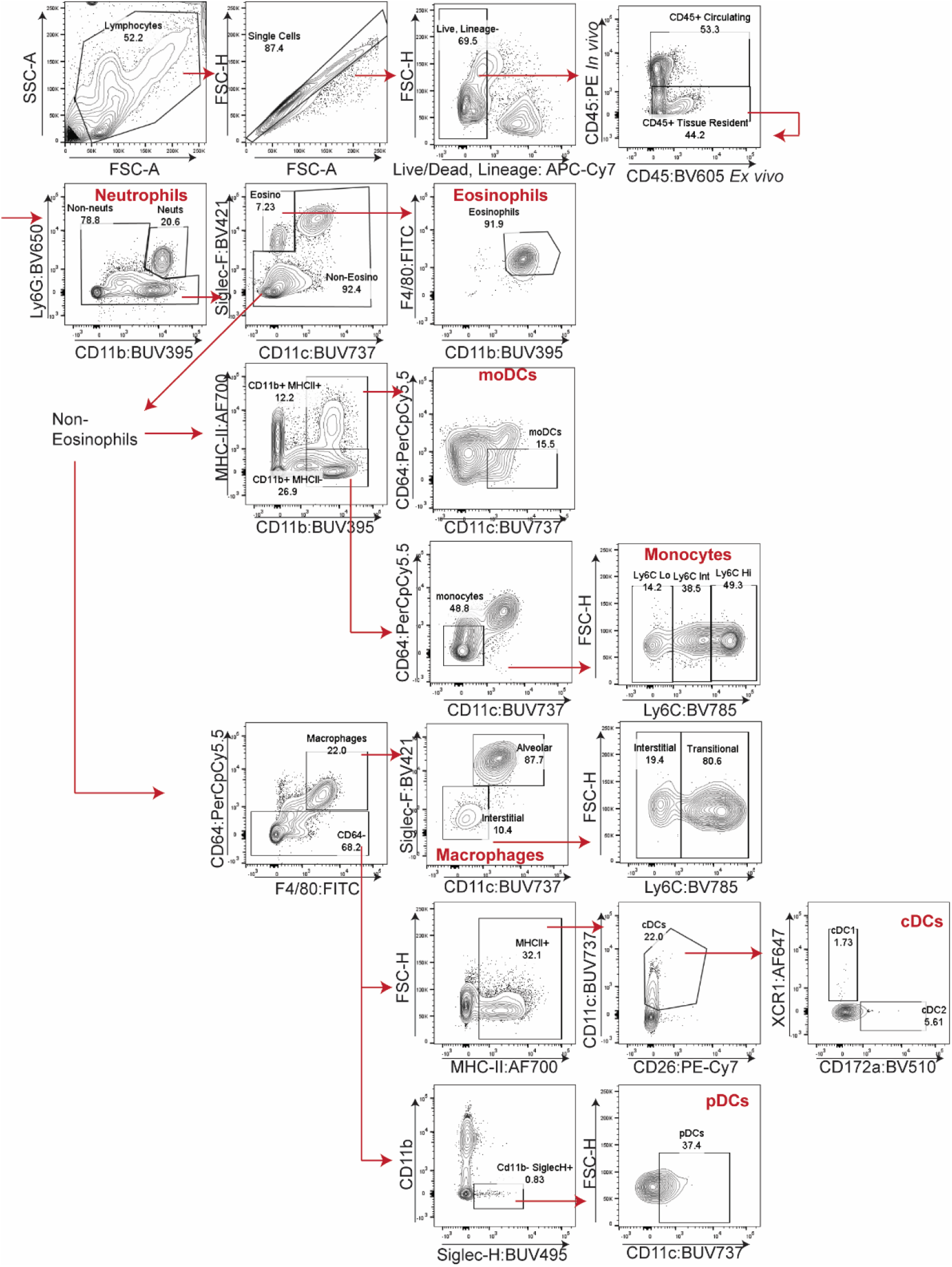
Gating strategy for innate immune populations in the mouse lung. C57Bl/6 mice were infected with MA-SARS-CoV-2 and lung tissue was harvested at 0, 2 and 4 days p.i. 5 minutes prior to euthanization, mice were intravitally labelled with CD45:PE (injected via the retro-orbital route). Lungs were processed to a single-cell suspension and analyzed via flow cytometry. Total cell populations were gated on (FSC-SSC), then singlets, then on Live (Ghost-Dye-) and Lineage- (CD19, CD3, NK1.1) populations. Lung infiltrating cells were identified as CD45 *ex vivo*+ and CD45 *in vivo*-. All following gates were made for both circulating and lung parenchymal populations. Neutrophils were identified as Ly6G+, CD11b+, CD11c-, and removed from downstream analysis. Eosinophils were identified as Siglec-F+, CD11c-, F4/80+, CD11b+, and removed from downstream analysis. Monocytes were identified as CD11b+, MHC-II-, CD11c-, CD64^low^. moDCs were identified as CD11b+, MHC-II+, CD64^low^, CD11c+. Macrophages were identified as CD64+, F4/80+, and distinguished into subsets based on Siglec-F, CD11c, and Ly6C expression. cDCs were identified as CD64-, MHC-II+, CD11c+, CD26+. cDC1s and cDC2s were distinguished by XCR1 and CD172a staining. pDCs were identified as CD11b-, Siglec-H+, and CD11c+. Subsequent gating is indicated by red arrows, and all final populations used for analyses are labelled in red. Flow plots are representative of lung populations from day 4 post MA-SARS-CoV-2 infection.

**Supplemental Figure 3.**
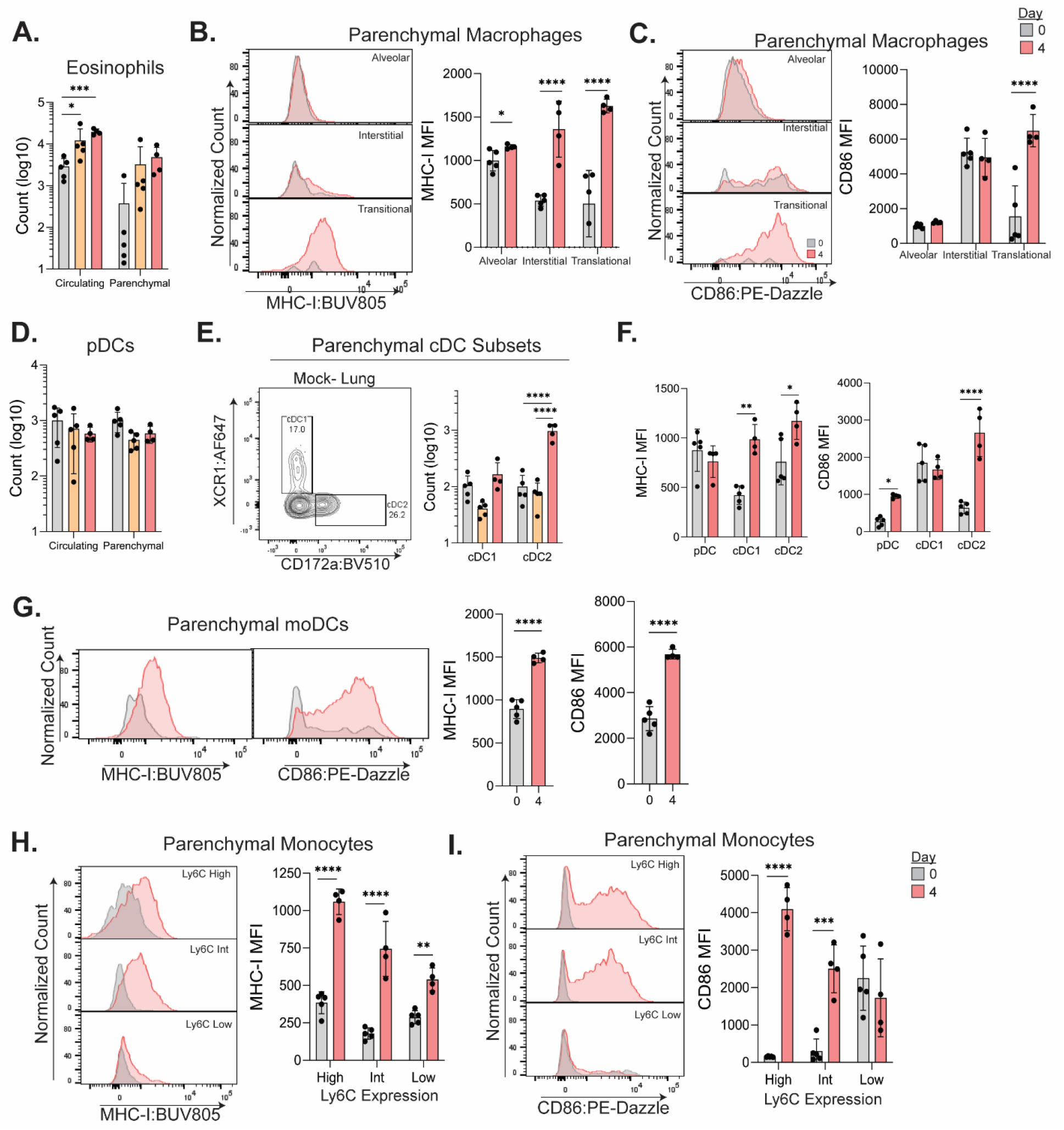
Lung parenchymal innate immune cells are activated in response to MA-SARS-CoV-2. C57Bl/6 mice were infected intranasally with MA-SARS-CoV-2, harvested at day 0, 2, or 4 p.i., and lung parenchymal vs circulating cells were identified as described in Fig. 2. A) Quantification of eosinophils (Lineage-, Ly6G-, CD11c-, SiglecF+, F4/80+, CD11b+). Representative histograms of B) MHC-I or C) CD86 expression on macrophage subsets. Quantified on the right are mean fluorescent intensities for day 0 and 4 p.i. D) Quantification of the number of pDCs (Lineage-, Ly6G-, CD64-, CD11b-, SiglecH+, CD11c+). E) Representative flow plot of cDC subset gating strategy and quantification of cDC1s (XCR1+, CD172a-) or cDC2s (XCR1-, CD172a+) on the right. F) Quantification of the MFI for MHC-I and CD86 on pDC and cDC subsets in the parenchyma at day 0 and 4 p.i. G) Representative histograms of MHC-I and CD86 expression on parenchymal moDCs at day 0 and 4 p.i., with the quantification on the right. Representative histograms for H) MHC-I or I) CD86 expression on lung parenchymal monocyte subsets and the corresponding quantification on the right. Results are representative of two independent experiments. Statistical significance was determined using unpaired one or two-way ANOVA. *p<0.05, **p<0.01, ***p<0.001, ****p<0.0001.

**Supplemental Figure 4.**
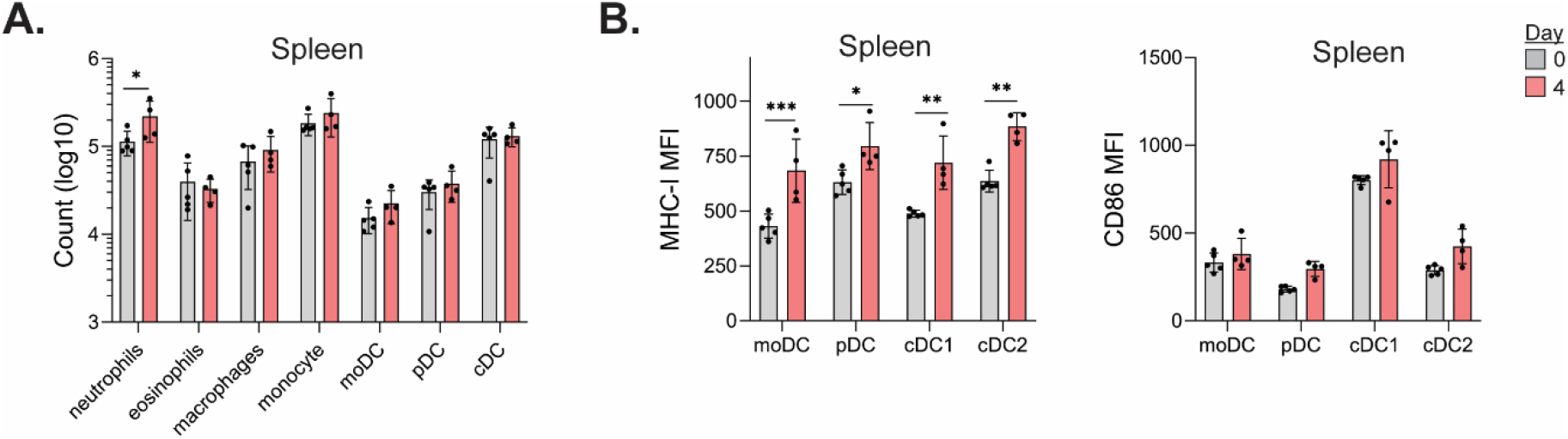
MA-SARS-CoV-2 infection induces systemic activation of dendritic cells. WT mice were infected with MA-SARS-CoV-2, and spleens were harvested at day 0 or 4 p.i. A) Quantification of the indicated cell populations in the spleen at day 0 or 4 p.i. D) MFIs for MHC-I (left) and CD86 (right) expressed on splenic dendritic cell populations. Results are representative of two independent experiments. Statistical significance was determined using unpaired one or two-way ANOVA. * = p<0.05, **= p<0.01, ***= p<0.001.

**Supplementary Figure 5.**
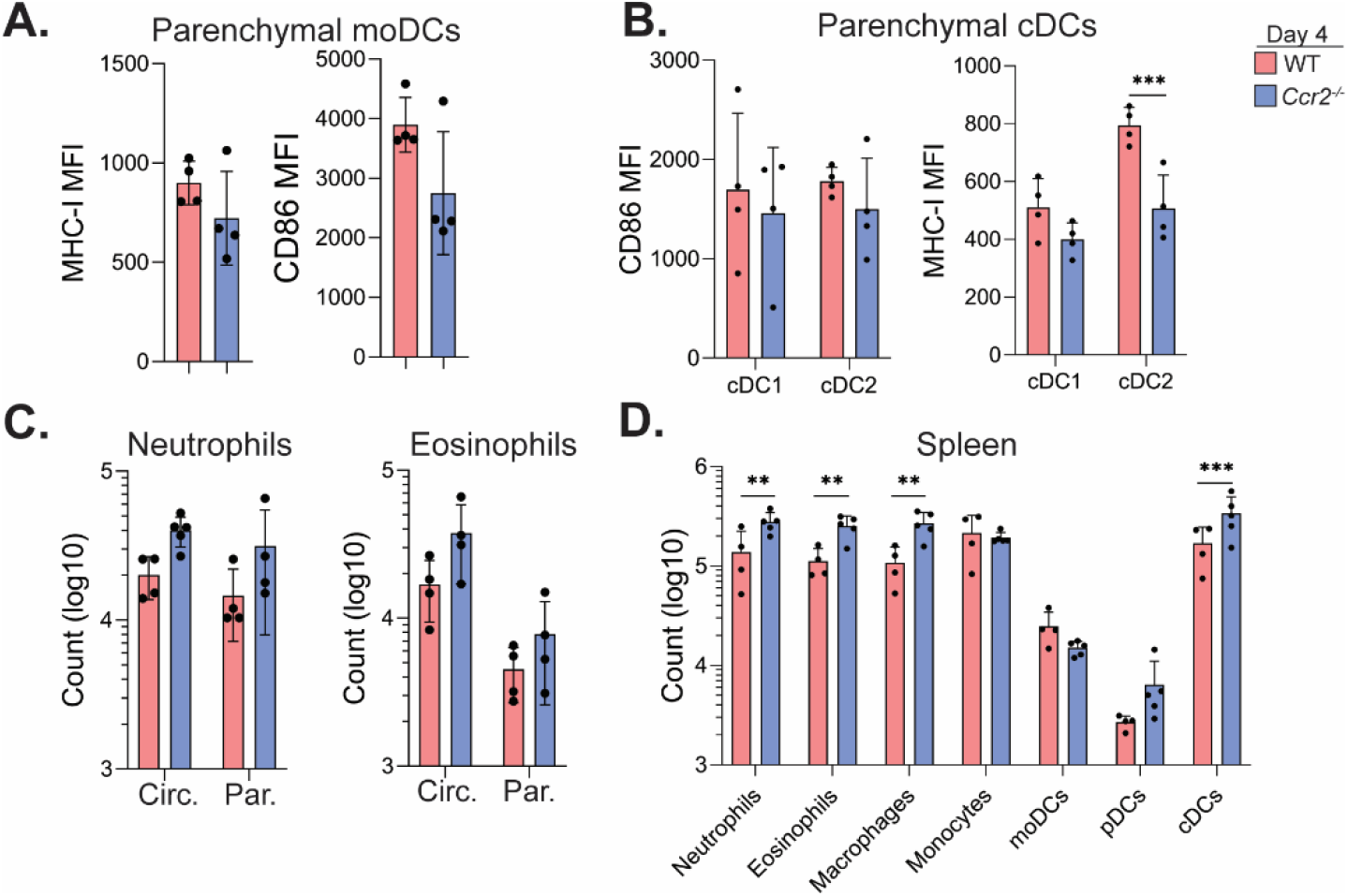
CCR2 limits systemic expansion of innate immune cells. WT and *Ccr2^-/-^* mice were infected with MA-SARS-CoV-2, and lung tissue and spleens were harvested at day 0 and 4 p.i. Lung-infiltrating cells were identified by intravital labelling and analyzed via flow cytometry. A) MFIs for MHC-I and CD86 expression on lung-infiltrating moDCs or B) cDC subsets. C) Quantification of the indicated granulocyte population in the lung. D) Quantification of the indicated innate immune cell population in the spleen. Results are representative of two independent experiments. Statistical significance was determined using unpaired student’s t-test or two-way ANOVA. * = p<0.05, **= p<0.01, ***= p<0.001.

## References

1 Zhou, P. et al. A pneumonia outbreak associated with a new coronavirus of probable bat origin. Nature 579, 270–273, doi:10.1038/s41586-020-2012-7 (2020).

2 Zhu, N. et al. A Novel Coronavirus from Patients with Pneumonia in China, 2019. N Engl J Med 382, 727–733, doi:10.1056/NEJMoa2001017 (2020).

3 Yin, X. et al. MDA5 Governs the Innate Immune Response to SARS-CoV-2 in Lung Epithelial Cells. Cell reports 34, 108628, doi:10.1016/j.celrep.2020.108628 (2021).

4 Arunachalam, P. S. et al. Systems biological assessment of immunity to mild versus severe COVID-19 infection in humans. Science 369, 1210–1220, doi:10.1126/science.abc6261 (2020).

5 Vanderheiden, A. et al. Type I and Type III Interferons Restrict SARS-CoV-2 Infection of Human Airway Epithelial Cultures. Journal of virology 94, doi:10.1128/jvi.00985-20 (2020).

6 Liao, M. et al. Single-cell landscape of bronchoalveolar immune cells in patients with COVID-19. Nature Medicine 26, 842–844, doi:10.1038/s41591-020-0901-9 (2020).

7 Wang, C. et al. Alveolar macrophage dysfunction and cytokine storm in the pathogenesis of two severe COVID-19 patients. EBioMedicine 57, doi:10.1016/j.ebiom.2020.102833 (2020).

8 Szabo, P. A. et al. Longitudinal profiling of respiratory and systemic immune responses reveals myeloid cell-driven lung inflammation in severe COVID-19. Immunity,doi:https://doi.org/10.1016/j.immuni.2021.03.005(2021).

9 Geissmann, F., Jung, S. & Littman, D. R. Blood monocytes consist of two principal subsets with distinct migratory properties. Immunity 19, 71–82, doi:10.1016/s1074-7613(03)00174-2 (2003).

10 Srivastava, M. et al. The Inflammatory versus Constitutive Trafficking of Mononuclear Phagocytes into the Alveolar Space of Mice Is Associated with Drastic Changes in Their Gene Expression Profiles. The Journal of Immunology 175, 1884, doi:10.4049/jimmunol.175.3.1884 (2005).

11 Lin, K. L., Suzuki, Y., Nakano, H., Ramsburg, E. & Gunn, M. D. CCR2+monocyte-derived dendritic cells and exudate macrophages produce influenza-induced pulmonary immune pathology and mortality. Journal of immunology (Baltimore, Md.: 1950) 180, 2562–2572, doi:10.4049/jimmunol.180.4.2562 (2008).

12 Seo, S. U. et al. Type I interferon signaling regulates Ly6C(hi) monocytes and neutrophils during acute viral pneumonia in mice. PLoS Pathog 7, e1001304, doi:10.1371/journal.ppat.1001304 (2011).

13 Cao, W. et al. Rapid differentiation of monocytes into type I IFN-producing myeloid dendritic cells as an antiviral strategy against influenza virus infection. Journal of immunology (Baltimore, Md. : 1950) 189, 2257–2265, doi:10.4049/jimmunol.1200168 (2012).

14 Xie, X. et al. An Infectious cDNA Clone of SARS-CoV-2. Cell Host Microbe 27, 841–848 e843, doi:10.1016/j.chom.2020.04.004 (2020).

15 Gu, H. et al. Adaptation of SARS-CoV-2 in BALB/c mice for testing vaccine efficacy. Science 369, 1603–1607, doi:10.1126/science.abc4730 (2020).

16 Hou, Y. J. et al. SARS-CoV-2 Reverse Genetics Reveals a Variable Infection Gradient in the Respiratory Tract. Cell, doi:https://doi.org/10.1016/j.cell.2020.05.042(2020).

17 Tegally, H. et al. Detection of a SARS-CoV-2 variant of concern in South Africa. Nature 592, 438–443, doi:10.1038/s41586-021-03402-9 (2021).

18 Guo, Q. et al. Induction of alarmin S100A8/A9 mediates activation of aberrant neutrophils in the pathogenesis of COVID-19. Cell Host & Microbe 29, 222–235.e224, doi:https://doi.org/10.1016/j.chom.2020.12.016(2021).

19 Tsou, C.-L. et al. Critical roles for CCR2 and MCP-3 in monocyte mobilization from bone marrow and recruitment to inflammatory sites. The Journal of Clinical Investigation 117, 902–909, doi:10.1172/JCI29919 (2007).

20 Chua, R. L. et al. COVID-19 severity correlates with airway epithelium–immune cell interactions identified by single-cell analysis. Nature Biotechnology 38, 970–979, doi:10.1038/s41587-020-0602-4 (2020).

21 Schulte-Schrepping, J. et al. Severe COVID-19 Is Marked by a Dysregulated Myeloid Cell Compartment. Cell 182, 1419–1440.e1423, doi:10.1016/j.cell.2020.08.001 (2020).

22 Silvin, A. et al. Elevated Calprotectin and Abnormal Myeloid Cell Subsets Discriminate Severe from Mild COVID-19. Cell 182, 1401–1418.e1418, doi:10.1016/j.cell.2020.08.002 (2020).

23 Xiong, Y. et al. Transcriptomic characteristics of bronchoalveolar lavage fluid and peripheral blood mononuclear cells in COVID-19 patients. Emerg Microbes Infect 9, 761–770, doi:10.1080/22221751.2020.1747363 (2020).

24 Fajnzylber, J. et al. SARS-CoV-2 viral load is associated with increased disease severity and mortality. Nature Communications 11, 5493, doi:10.1038/s41467-020-19057-5 (2020).

25 Peters, W. et al. Chemokine receptor 2 serves an early and essential role in resistance to Mycobacterium tuberculosis. Proc Natl Acad Sci U S A 98, 7958–7963, doi:10.1073/pnas.131207398 (2001).

26 Roy, R. M., Wüthrich, M. & Klein, B. S. J. T. J. o. I. Chitin elicits CCL2 from airway epithelial cells and induces CCR2-dependent innate allergic inflammation in the lung. 189, 2545–2552 (2012).

27 Liegeois, M., Legrand, C., Desmet, C. J., Marichal, T. & Bureau, F. The interstitial macrophage: A long-neglected piece in the puzzle of lung immunity. Cellular immunology 330, 91–96, doi:10.1016/j.cellimm.2018.02.001 (2018).

28 Guilliams, M. et al. Alveolar macrophages develop from fetal monocytes that differentiate into long-lived cells in the first week of life via GM-CSF. The Journal of experimental medicine 210, 1977–1992, doi:10.1084/jem.20131199 (2013).

29 Hashimoto, D. et al. Tissue-resident macrophages self-maintain locally throughout adult life with minimal contribution from circulating monocytes. Immunity 38, 792–804, doi:10.1016/j.immuni.2013.04.004 (2013).

30 Yona, S. et al. Fate mapping reveals origins and dynamics of monocytes and tissue macrophages under homeostasis. Immunity 38, 79–91, doi:10.1016/j.immuni.2012.12.001 (2013).

31 Nakano, H., Lyons-Cohen, M. R., Whitehead, G. S., Nakano, K. & Cook, D. N. Distinct functions of CXCR4, CCR2, and CX3CR1 direct dendritic cell precursors from the bone marrow to the lung. J Leukoc Biol 101, 1143–1153, doi:10.1189/jlb.1A0616-285R (2017).

32 Herold, S. et al. Lung epithelial apoptosis in influenza virus pneumonia: the role of macrophage-expressed TNF-related apoptosis-inducing ligand. The Journal of experimental medicine 205, 3065–3077, doi:10.1084/jem.20080201 (2008).

33 Grant, R. A. et al. Circuits between infected macrophages and T cells in SARS-CoV-2 pneumonia. Nature, doi:10.1038/s41586-020-03148-w (2021).

34 Chen, R. E. et al. Resistance of SARS-CoV-2 variants to neutralization by monoclonal and serum-derived polyclonal antibodies. Nat Med, doi:10.1038/s41591-021-01294-w (2021).

